# Epigenetic information loss is a common feature of multiple diseases and aging

**DOI:** 10.1101/2023.05.07.539727

**Authors:** Naor Sagy, Chieh Chang, Maayan Gal, Daniel Z Bar

**Affiliations:** The faculty of Medicine, Tel Aviv University, ISRAEL

## Abstract

Aging is a major risk factor for a plethora of diseases. The information theory of aging posits that epigenetic information loss is a principal driver of the aging process. Despite this, the connection between epigenetic information loss and disease has not been thoroughly investigated. Here, we analyzed tissue-unique methylation patterns in healthy and diseased organs, revealing that for several diseases these patterns degrade, regressing to a mean form. We interpret this as epigenetic information loss, where tissue-unique patterns erode. Information loss is not limited to diseases. Age-related erosion of unique methylation patterns was observed in some tissues and cells, while other tissues and cells diverged away from the mean. Our findings demonstrate that analyzing methylation patterns in tissue-unique sites can effectively distinguish between patients and healthy controls across a range of diseases, and underscore the role of epigenetic information loss as a common feature in various pathological conditions.

Graphical abstract
Tissue unique methylation pattern regress toward the mean upon disease.
A single methylation site, showing low methylation in the liver and high in every other tissue, becomes more methylated in diseased livers.

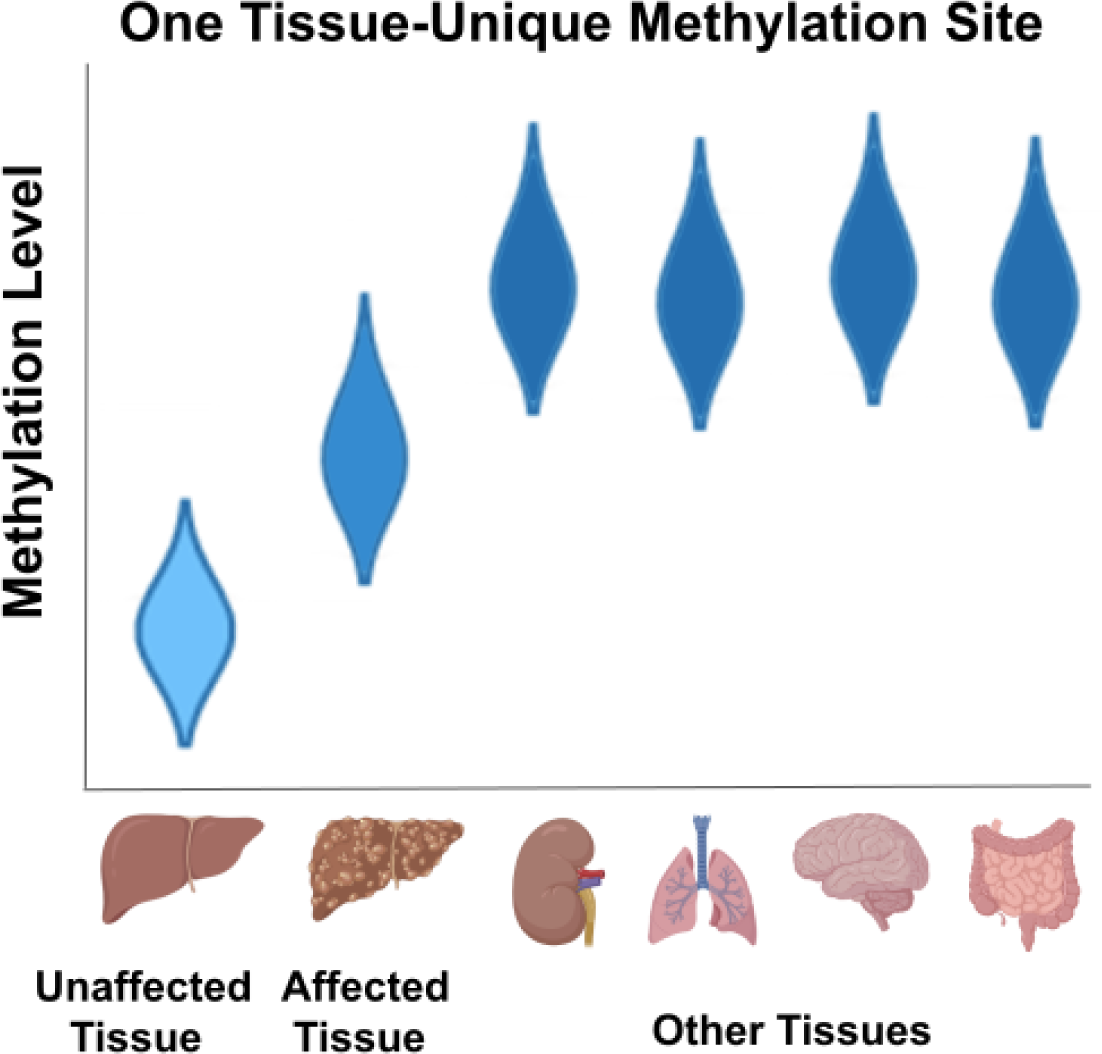

## Introduction

Aging is the primary risk factor for numerous chronic diseases, such as cardiovascular diseases, cancer, and type 2 diabetes (T2D) ^1^. Epigenetic alterations, including changes in methylation levels at specific genomic loci, are a well-known hallmark of aging ^2–4^. Linear combinations of these loci, known as epigenetic clocks, strongly correlate with chronological age ^5^ and can predict all-cause mortality ^6^. Behaviors that correlate with decreased healthspan and lifespan, including smoking and bad diet, accelerate some epigenetic clocks ^6^. Stress may also induce rapid increase in biological age, as measured by epigenetic clocks ^7^. Interestingly, recovery from stress may revert these changes ^7^. Similarly, heterochronic parabiosis reduced the epigenetic age ^8^. However, the causality of these methylation changes and why certain loci correlate with age remain unresolved ^2^.

The epigenetic theory of aging posits that the loss of epigenetic information disrupts youthful gene expression patterns, leading to cellular dysfunction and senescence ^9–12^. Thus, information loss is suggested to drive the aging process ^13^. Age-dependent loss of youthful gene expression patterns has been demonstrated in some cases ^14–16^, but not in others ^17^. By contrast, single-cell level analysis of global coordination also suggests that transcriptional dysregulation is common or universal with aging ^18^.

Medical diagnosis typically relies on prior knowledge and the identification of relevant pathologies to explain symptoms. Specific DNA methylation patterns strongly correlate with medical conditions, but these correlations are only inferred by comparing methylation patterns of labeled cohorts ^19–22^. In this study, we demonstrate that tissue-specific methylation patterns change with age and the progression of certain diseases. This pattern change allows us to predict Chronic Kidney Disease (CKD), liver diseases, T2D, and sun exposure in healthy skin using methylation data alone, without any additional information or prior knowledge. Moreover, in multiple cases, the direction of change is not random; instead, it moves towards a common form across all other tissues, which we interpret as epigenetic information loss. Our findings establish the loss of epigenetic information as a shared feature of multiple diseases.

## Results

### Identification of tissue-specific methylation sites

Epigenetic information loss can take many forms, and can be challenging to identify and quantify in a consistent manner across multiple diseases, tissues and datasets. It was previously shown that in the kidney, methylation sites with a kidney-unique signature always regress towards the common form with CKD progression ^23^. The unique methylation signature of the kidney, as manifested in these sites, erodes as the tissue becomes more similar to the rest of the body. The existence of tissue-unique methylation sites allows the exploration of such information loss, in a manner that is both tissue-specific and consistent across all examined tissues.

To facilitate this analysis, we generated a dataset of all tissue-specific methylation sites (**Fig. 1A-C**) based on the NGDC-CNCB dataset ^24^. Of the 30 tissues across 5323 samples in the dataset, 23 had unique methylation sites, ranging from 55 in bladder to 50,872 in placenta (**Fig. 1D**; **Supp. Table 1**). In total, 87,922, or ∼18% of sites tested, displayed a unique methylation pattern in at least one tissue, with over half of them in the placenta.

**Figure 1:**
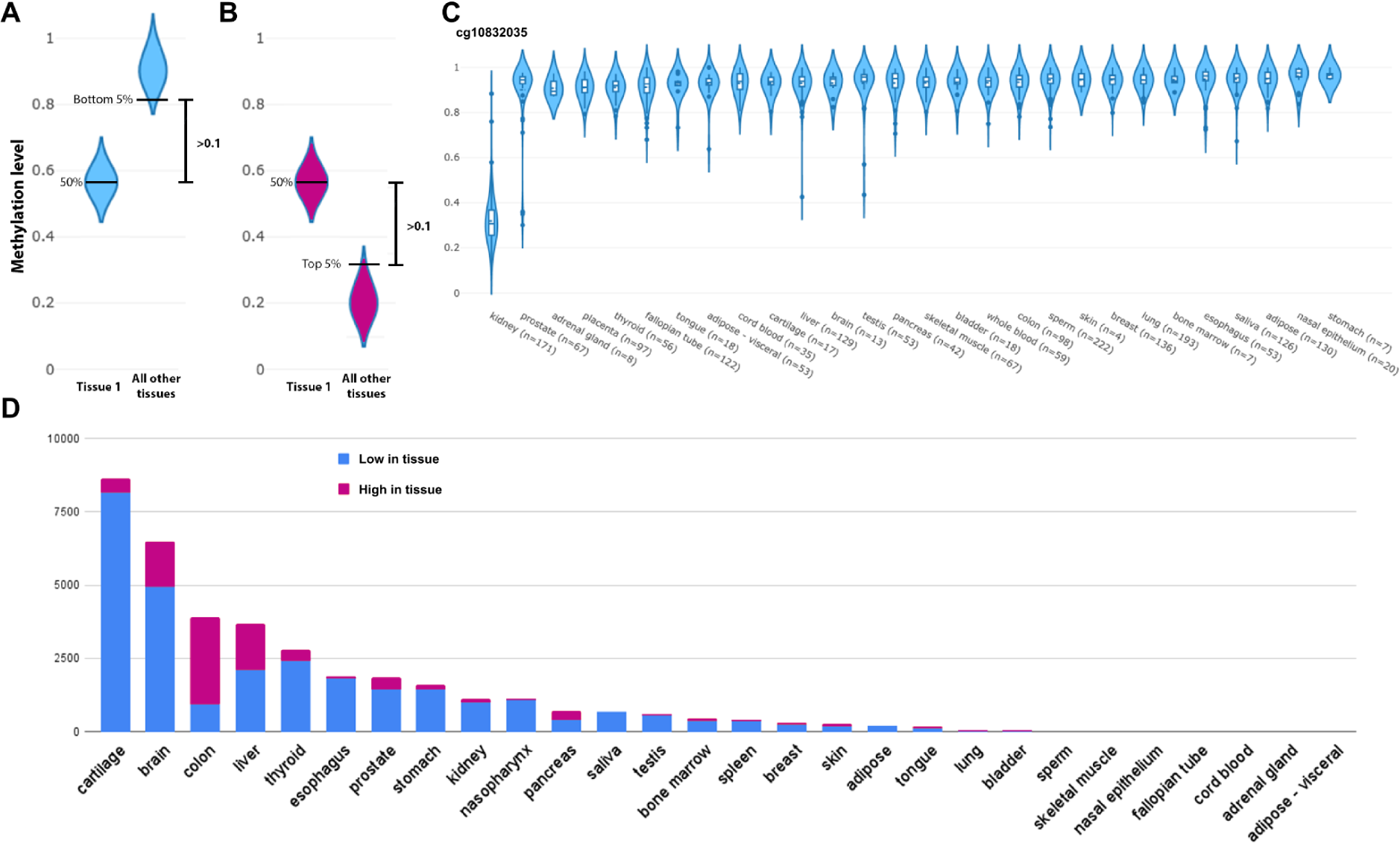
Identification of tissue-specific methylation sites. Methylation sites that show a median absolute methylation value at least 0.1 below the bottom 5% quantile (**A**) or above the top 5% quantile (**B**) are considered unique. For example (**C**) shows a unique methylation pattern in the kidney. (**D**) Unique methylation sites for each of the tissues, excluding the placenta, that had over 40,000 low and 10,000 high sites.

### Hypothesis-free mapping of kidney-unique methylation sites correlates with CKD

Epigenetic information loss occurs in nearly all uniquely methylated sites as CKD progresses (^23^ and **Supp. Tables 2-3**). To obtain an unbiased, global view of these data, we projected the methylation values of kidney sites (**Supp. Tables 1-3**) into a 2D space using principal component analysis (PCA). This mapping spatially separated individuals with high and low interstitial fibrosis (**Fig. 2A**). To determine if this is a general feature across all methylation sites, we repeated the analysis with randomly selected methylation sites (**Fig. 2B**) and extended the controls to sites with high (**Fig. 2C**) and low (**Fig. 2D**) kidney methylation variability across individuals. Among all cases, uniquely methylated sites demonstrated the strongest spatial separation. We concluded that uniquely methylated sites can be utilized for unbiased characterization of kidney functional decline.

**Figure 2:**
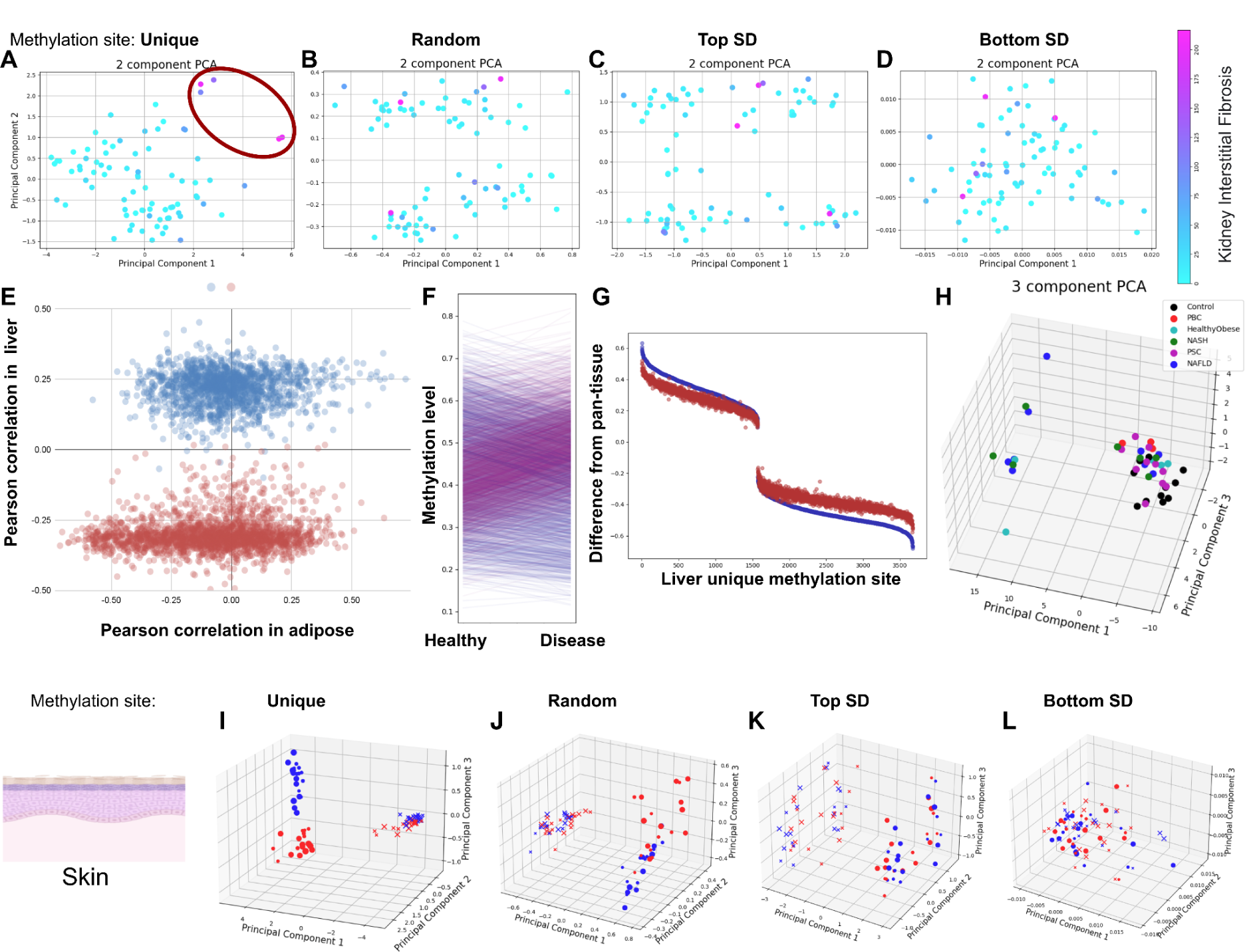
Epigenetic information loss in disease and stress. PCA analysis of kidney-unique methylation sites clusters kidneys with high levels of interstitial fibrosis (**A**), but not when random (**B**), high variability (**C**) or low variability (**D**) sites are used. (**E**) Liver-unique methylation sites show a binomial distribution of correlations to liver diseases in liver samples (Y axis), but not in adipose (X axis). More-methylated samples (red) show a negative correlation, while less-methylated samples (blue) show a positive correlation. (**F**) Methylation levels (mean) for sites that are more (red) and less (blue) methylated in the liver (compared to the rest of the body) reveals regression to the mean in disease. (**G**) Differences between each liver-unique methylation site mean and pan-tissue average, in healthy (blue) and sick (red) samples. Regression to the mean (red closer to zero) is observed in nearly all sites. (**H**) PCA of liver-unique methylation sites in men only. Separation based on tissue-unique sites following environmental stress: 3D PCA analysis of skin-unique methylation sites separated sun-exposed (red) from sun-protected (blue) samples (**I**), but not when random (**J**), high variability (**K**) or low variability (**L**) sites are used. X - dermis; o - epidermis; size indicates young (small) or old (large).

### Multiple liver diseases show loss of epigenetic information

The liver is renowned for its exceptional regeneration capacity, which may make it less susceptible to aging effects compared to other tissues. Indeed, the risk of some liver diseases does not appear to correlate with age, at least in men ^25,26^. However, liver function declines with age, and aging is associated with increased severity and poor prognosis of multiple liver diseases ^27^. To explore the role of epigenetic information loss in liver diseases, we analyzed a combined multi-disease and multi-tissue dataset (^28^; GSE61256; N = 137; **Supp. Tables 2-3**). It includes, in addition to liver methylation samples from healthy individuals, data for healthy but obese participants, as well as Primary Biliary Cirrhosis (PBC), Non-Alcoholic SteatoHepatitis (NASH), Primary Sclerosing Cholangitis (PSC) and Non-Alcoholic Fatty Liver Disease (NAFLD) patients. Methylation data for adipose and muscle tissues were also available for some donors. Our tissue-specific dataset contains 3,673 liver-specific methylation sites, all of which were found in this dataset, enabling high-confidence detection of epigenetic information loss. The correlation between liver methylation in the NGDC-CNCB dataset, used to derive the unique sites, and the controls of GSE61256, was r = 0.99 (**Supp. Fig. 1A**), indicating high technical reproducibility across different datasets. We first explored whether methylation levels in these sites correlate with all diseases. Remarkably, liver-unique methylation sites exhibited a bimodal distribution with a positive and a negative peak, and almost no sites showed low correlation to liver disease (**Fig. 2E**, Y-axis). In contrast, the same sites in the same patients, but in adipose tissue, showed a normal distribution (**Fig. 2E**, X-axis). We investigated what distinguishes sites that positively vs. negatively correlate with liver disease. Overwhelmingly, these two populations were distinguished by their methylation status in the liver compared to the rest of the body (3,658 of 3,673; p < 10^-1000^). Sites that displayed lower methylation in the liver, as compared to the rest of the tissues (blue), had higher methylation values in diseased livers (positive correlation), while sites with higher liver methylation (red) had decreased methylation in diseased livers (negative correlation). Thus, 99.6% of these sites regressed to the mean with liver disease progression (**Fig. 2F-G**).

Contrary to our expectations based on these results, PCA of these samples showed a poor separation between healthy and diseased livers (**Supp. Fig. 1B**). We hypothesized that another source of variance was dominating this analysis. Indeed, we found that men and women formed two distinct clusters when performing PCA using high-variability sites (**Supp. Fig. 1C**), in line with previous findings that the human liver has a sex-specific methylation profile in autosomes ^29^. When analyzing men separately, controls clustered together and partially separated from diseases (**Fig. 2H**). Control methylation sites exhibited weaker separation of the control group (**Supp. Fig. 1D**). Surprisingly, analyzing women only yielded a relatively poor separation of controls (**Supp. Fig. 1E**). That can be attributed to gene expression differences between men and women ^30–32^. These inherent dissimilarities might manifest in methylation profiles and different hormone levels in the liver ^33^, and the nature of PCA analysis which segregates samples according to maximum variance and not by a certain biological orientation. These results suggest that liver disease results in epigenetic information loss in the liver.

### Type 2 diabetes shows epigenetic information loss in adipose but not in the pancreas

T2D is the most prevalent form of diabetes, characterized by high blood sugar, insulin resistance, a relative lack of insulin, and changes to the pancreas ^34–36^. Age is a major risk factor for T2D, and it correlates with increased biological age as indicated by epigenetic clocks ^37^. To investigate if epigenetic information is lost in the pancreas of T2D patients, we applied our analysis to a dataset mapping 27,578 CpG sites in human pancreas samples from T2D patients and controls (^38^; GSE21232; N = 16). Of the 705 pancreas-unique methylation sites, only 45 were covered by the methylation array used. Among these, 28 were less methylated in the pancreas. Contrary to our findings in liver and kidney, 26 of the 28 sites diverged away from the mean in T2D (**Fig. 3A**; p < 10^-5^). Of the 17 sites that were more methylated in the pancreas, divergence was observed in 16 (**Fig. 3A**; p < 10^-3^). PCA almost completely separated diabetic from non-diabetic pancreas samples (**Supp. Fig. 2A**). Interestingly, using random sites or sites with high variance in the pancreas — but not sites of low variance — also separated diabetic from non-diabetic pancreases (**Supp. Fig. 2B**). Thus, the epigenetic changes are robust enough for T2D to be detected in multiple sites.

**Figure 3:**
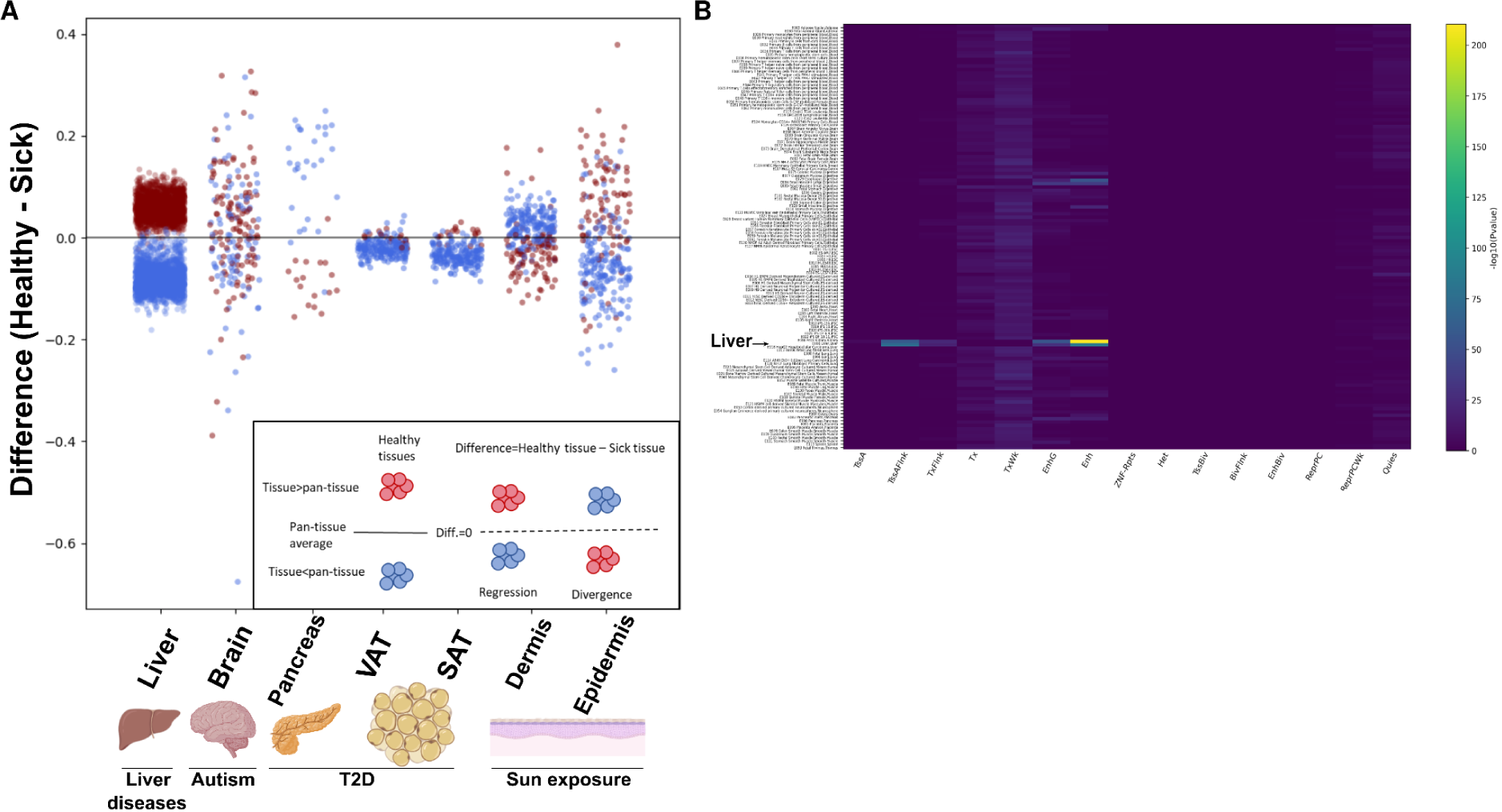
Methylation difference in disease across seven tissues. **(A)** Inset - schematic explanation: Red dots represent sites methylated above other tissues, while blue dots represent lower methylation levels. When looking at the difference between healthy and sick, red above zero and blue below indicate regression to the mean, while the opposite indicates divergence. Y-axis: Difference in methylation levels, for each tissue-unique methylation site, between healthy and sick (cohort averages). Differences shown for liver (liver diseases), Brain (autism), pancreas (T2D), VAT and SAT (T2D) and dermis and epidermis (sun exposure). (**B**) Enrichment of liver unique (low in liver) methylation sites across 15 chromatin states (x-axis) and 127 tissues and cell lines (y-axis). Strongest enrichment, highlighted with “liver” text and arrow, was observed for methylation in liver tissue (E066), followed by hepatocellular carcinoma (E118).

Diabetes is closely linked to obesity and the accumulation of adipose tissue. However, not all adipose tissues confer the same risk. Specifically, visceral adipose (VAT) poses a significantly greater risk than subcutaneous fat (SAT) ^39^. To test if epigenetic information is lost in diabetes, we examined a dataset of VAT and SAT from T2D patients and controls (^40^; GSE162166; N = 40). Globally, some methylation patterns were observed. In VAT, 84% of all methylation sites showed elevated methylation in T2D patients. Furthermore, examining methylation sites with a robust change in any direction—measured by an absolute correlation greater than 0.3—94% of sites showed increased methylation. Interestingly, this was observed in VAT only, whereas in SAT, only 46% showed elevated methylation. Adipose tissue has 222 unique sites (**Fig. 1D**, **Supp. Tables 2-3**). Among these, 212 were covered by the methylation profiling dataset, including VAT and SAT from T2D patients and controls (^40^; GSE162166). Methylation patterns regressed towards the mean at 90% of these sites in VAT and 95% in SAT (**Fig. 3A** - blue dots below the zero line and red dots above it). Although statistically significant, this finding is not particularly surprising, as 91% of the unique sites exhibited lower methylation levels in adipose tissue, implying that the general trend in VAT can largely account for this observation. Considering the remaining 9% (N = 18) of the sites, regression towards the mean (and contrary to the 84% trend observed) was evident in 9 of them (p = 10^-3^). In SAT, 14 of the 18 sites regressed towards the mean (p < 10^-7^). Consequently, we concluded that a global increase in methylation is a characteristic feature of VAT, but not SAT, in T2D patients, and both tissues experience a loss of epigenetic information. However, these findings were not accompanied by clear separation by PCA, as healthy and T2D patients did not form distinct clusters in neither VAT nor SAT, when using adipose-unique sites (**Supp. Fig. 2C**).

### Autism does not show significant epigenetic information loss

Autism is a neurodevelopmental disorder characterized by difficulties with social interaction and communication. Symptoms of autism typically appear early in childhood, and it has not been linked to aging. We tested if epigenetic information is lost in autism. The brain contains a relatively large number of unique methylation sites: 6,491. Of these, 209 were covered by a dataset of brain methylation (^41^; GSE38608; N = 36). In autism, we observed information loss in 125 of these. While these findings were marginally statistically significant (p = 0.005), they appeared very weak compared to kidney and liver diseases (**Fig. 3A** - no clear separation of red and blue above or below the zero line in brain, unlike in the liver). We performed PCA using all brain-unique methylation sites, and autistic brains appeared indistinguishable from controls (**Supp. Fig. 2D**). A clear signature was observed, but it was not related to autism (see “Epigenetic information loss during aging” below). Together, these results suggest that loss of tissue-specific methylation information is not a universal feature of all diseases and disorders. However, these results do not exclude the possibility that a different analysis or sample correction may reveal information loss.

### Epigenetic information is lost in environmentally stressed tissue

The skin serves as a barrier that protects the body from external insults, such as UV radiation from the sun. Sun exposure prematurely ages the skin, termed photoaging ^42^. Photoaging manifests differently in the dermis and epidermis, which can be partly attributed to the limited penetration of UV-B into the dermis ^43^. We investigated whether typical environmental stressors to tissue induce epigenetic information loss. To do so, we compared methylation data from sun-exposed and sun-protected skin samples of both dermis and epidermis, in young and old individuals ^44^. Of the 277 skin unique methylation sites, 275 were present on the dataset (GSE51954; N = 78). In the dermis, divergence from the mean upon sun exposure was observed (**Fig. 3A** - dermis. Blue dots above zero, red below; **Supp. Table 3**). In contrast, in the epidermis, regression to the mean occurred at 223 and 226 sites for young and old individuals, respectively (**Fig. 3A**; **Supp. Table 3**). When mapping methylation-levels of skin-unique methylation sites into 3D space using PCA, dermis and epidermis samples were clearly separated (**Fig. 2I**). However, samples from older versus younger donors exhibited only partial separation (**Fig. 2I**). Sun-exposed and sun-protected samples from each tissue type clustered separately, with the expected stronger separation observed in the epidermis. Analyzing the same samples using methylation sites that were not tissue-specific, including random sites and sites with high standard deviation, did not yield such a distinct separation (**Fig. 2J-L**). Consequently, we conclude that sun exposure is associated with epigenetic information loss in the epidermis.

### Epigenetic information loss during aging

The epigenetic theory of aging posits that the loss of epigenetic information contributes to the aging process (^10,13,45,46^). Tissue functionality declines with age, similar to the deterioration observed in multiple diseases. This decline may vary among tissues, as some experience a faster loss of function on average than others. Nonetheless, a strong epigenetic signature of aging has been observed in most tissues ^5^. We tested if tissue-unique epigenetic information is lost during aging, focusing on three tissues. The brain has low regenerative capacity and slow cell turnover. With aging, cognitive functions decline and memory formation may become impaired. In contrast, the intestine exhibits high regenerative capacity and cell turnover. Aging in the intestine is characterized by reduced self-renewal capacity of intestinal stem cells and diminished tissue self-repair functionality ^47^. When plotting the brain-unique methylation sites for autistic and nonautistic individuals, we noticed PCA separated the samples by age (**Supp. Fig. 2E** and **Fig. 4A**). Indeed, PC1 strongly correlated with the age of the individuals (**Fig. 4A**). However, random sites, as well as sites with high variance also produced the same result (**Fig. 4B-D**). Of the 209 brain-unique sites covered by the dataset, regression to the mean was observed in 148, and not observed in 61 sites (p < 10^-9^). Next we analyzed normal colon mucosa samples (^48^; GSE48988; N = 178). Of the 3894 intestine-unique sites, 290 were found in the dataset. Of these, 87% (252) had higher methylation levels in the intestine than in other tissues. 48 sites did not produce any methylation values and were discarded. Divergence from the mean was observed in 237 of the 242 remaining sites, while regression was observed in just 5 sites (p < 10^-63^; **Fig. 4J**). PCA generated a clear age gradient when using intestine-unique sites, but not in any of the controls (**Fig. 4F-I**). We concluded that the tissue-unique epigenetic signature degrades with age, but only in some tissues. This result is independent of whether unbiased PCA clustering separates samples by age.

**Figure 4:**
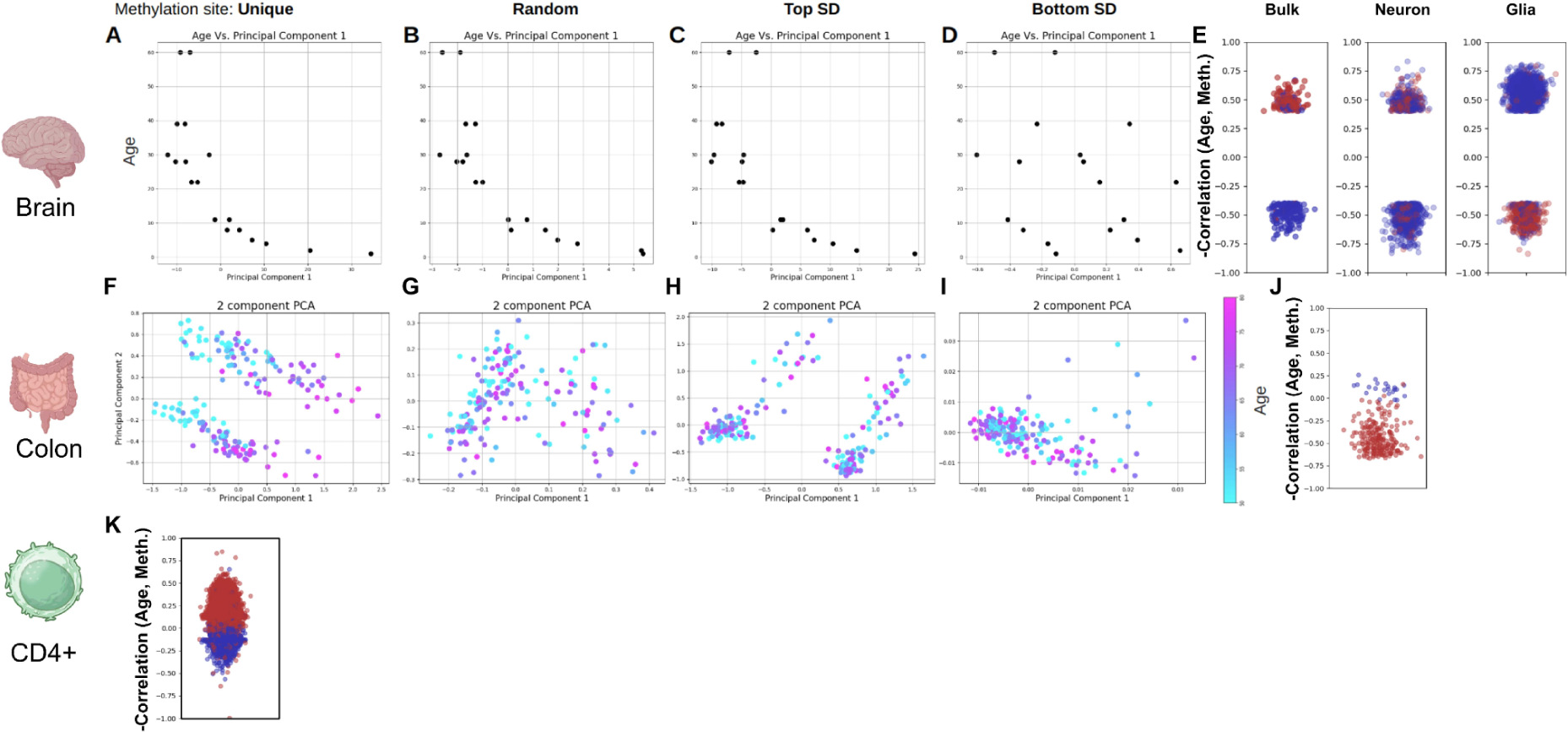
Epigenetic information loss in aging. PC1 vs age in brain samples, using (**A**) unique (**B**) random (**C**) high variability and (**D**) low variability sites. (**E**) Correlation of methylation levels and age for brain-unique methylation site, in bulk, neuronal and glial cells. PCA of intestine samples, using (**F**) unique (**G**) random (**H**) high variability and (**I**) low variability sites. Color indicates age.(**J**) Correlation of methylation levels and age for intestine-unique methylation site (**K**) Correlation of methylation levels and age for intestine-unique methylation site. All correlations are plotted inversely for schematic consistency (red above zero for regression).

### Functional analysis of tissue-unique methylation sites

Assigning functional significance to the methylation sites underlying epigenetic clocks remains challenging ^49^. We have previously shown that kidney-unique methylation sites are enriched for disease relevant transcription factors and the less-methylated sites are enriched for kidney tissue-specific enhancers ^23^. To test if this is a general attribute of tissue-unique methylation sites, we selected from the 30 tissues analyzed for unique methylation sites (**Fig. 1D**; **Supp. Table 1**) and identified the tissues most similar to the seven disease tissues investigated (**Fig. 3A**). The low-in-tissue (blue) methylation sites were analyzed using eFORGE 2 for enrichment of 15 chromatin marks across and multiple samples ^50^. In all tissues analyzed, the strongest enrichment was observed for tissue-specific enhancers of tissues most proximal to the analyzed tissue (**Fig. 3B, Supp. Fig. 3A** ^51^). For example, sites showing low methylation in liver and high in all other tissues were most enriched for enhancers identified in liver tissue (**Fig. 3B**). The only exception to this was in brain tissue, where strong enhancer enrichment was additionally observed in ES-derived ectoderm CD56+ cell culture, the primary germ layer of origin for most brian cells, with a marker crucial for neuronal development, as well as ES-derived neuronal progenitor cells (**Supp. Fig. 3A**). We concluded that unique undermethylated sites are enriched for tissue-specific enhancers, and these get more methylated in some age-associated diseases. Additionally, tissue-unique methylation sites showed enrichment for known transcription-factor binding sites, with limited overlap between motifs identified in different tissues (**Supp. Fig. 3B**).

### Epigenetic information is lost in some, but not all, cell types

To investigate whether epigenetic information loss occurs within a cell population, we analyzed a dataset of human brain samples (^52^; GSE41826; N = 145) which included bulk tissue samples as well as fluorescence-activated cell sorted (FACS) neuronal and non-neuronal (predominantly glia) cells. In the bulk samples, consistent with our previous observation, regression to the mean with age was observed in 85% (1302/1531; p < 10^-181^) of more methylated sites and 77% (3819/4926; p < 10^-344^) of less methylated sites. Analyzing only sites that robustly correlate with age (R > 0.4) increased these numbers to 96% (p < 10^-31^; **Fig. 4E**) and 86% (p < 10^-30^) respectively. This trend was mimicked in neurons, where regression to the mean was observed in 71% (p < 10^-9^) and 70% (p < 10^-32^), of more and less methylated sites, respectively. Surprisingly, the opposite trend was observed in glia cells, where methylation levels predominantly diverged away from the mean (85%; p < 10^-49^ and 94%; p < 10^-390^, respectively).

To further corroborate these findings, we analyzed the NGDC-CNCB blood DNA methylation dataset ^53^. This independent dataset comprises 25 types of blood samples (N=3402), including mixed and cell-type-specific samples. We focused on purified naive CD4+ T cells (CD4+). As these cells were not present in our original analysis, we identified 29,137 methylation sites unique to CD4+ cells (**Supp. Table 4**), compared to the remaining 24 blood sample types. Correlation of methylation level and age was calculated for each unique site for samples where age was available (N=71). The majority of the sites were more methylated in CD4+ compared to the remaining samples. Of these, 96% (20825/21556; p < 10^-1000^) of sites regressed towards the mean with age (**Fig. 4K** - red dots). Similarly, of the less methylated sites 93% (7102/7581; p < 10^-1000^) regressed towards the mean (**Fig. 4K** - blue dots). Thus, some cell types regress to the mean, while others diverge away from it.

### Methylation-based disease classification

Tissue-specific methylation signature changes in various diseases raise the question of whether these unique sites can be used for disease classification. We applied three classification models using only methylation data. We were able to distinguish between low and high GFR (kidney) with an accuracy as high as 82.35%, between sun-exposed and sun-protected skin samples in the dermis with an accuracy of 80%, and in the epidermis with an accuracy of 100%, respectively (**Supp. Fig. 4**)

## Discussion

Aging is accompanied by epigenetic changes. By focusing on DNA methylation sites with unique signatures compared to other tissues, we could isolate genomic loci where age-dependent changes in methylation can be interpreted. In these sites, observing the direction of change can inform us if it is random, hypomethylation, hypermethylation, divergence or regression to the mean. In multiple instances, including diseases, environmental stress, and natural aging, these sites predominantly changed by regression to the mean. For example, 99.6% of liver-unique sites regressed to the mean in liver diseases. We interpret these results as a loss of epigenetic information. The unique methylation signature that characterizes the tissue becomes more similar to other tissues in the body, akin to the smoothening of the epigenetic landscape or exdifferentiation reported elsewhere ^11,13,54^. These results do not imply that a similar process does not occur in other methylation sites. On the contrary, the fact that such a large fraction of the a-priori selected sites showed this trend suggests this is likely to be a more general phenomenon. Moreover, other types of epigenetic information may exhibit a similar phenomenon. However, analyzing these sites will require the development of expectations as to the direction of change, to distinguish information loss from changes that may reflect different processes. Interestingly, some tissues that do not show this phenomenon demonstrate the opposite effect. Methylation sites diverge away from the common form as the tissue becomes more distinct. Thus, our experiments do not support a role of epigenetic information loss, at least for these sites and bulk tissue. We postulate that another unknown mechanism, actively counteracting the smoothening of the epigenetic landscape, is dominant in these samples.

We found that tissue unique methylation sites are enriched for enhancers specific to the same tissue (**Fig. 3B** and **Supp. Fig. 3A**). These sites tend to become more methylated, and thus typically less accessible, with the progression of some age-associated diseases (**Fig. 3A)**. One could speculate that this closure of tissue-specific enhancers leads to loss of tissue-specific chromatin architecture and drives the smoothening of the epigenetic landscape. However, additional data is needed to infer causality and substantiate such a speculation.

One possible explanation for the loss of epigenetic information in disease and aging is changes to tissue cell composition. Such changes, including replacement of resident-tissue cells by fibroblasts and accumulation of immune cells, have been demonstrated in aging and disease in some tissues, but not in others ^55–58^. A non-exclusive alternative is changes within the cell population. For example, in epithelial to mesenchymal transition (EMT), epithelial cells undergo changes that enable them to assume a mesenchymal cell phenotype ^59^. EMT has been associated with fibrosis in kidney, liver, and intestine ^59–63^ and changes in DNA methylation have been suggested to causally underlie EMT ^64^. In human brain samples, regression to the mean with age was observed in brain-unique sites of bulk tissue. When looking at sorted cells, we observed regression to the mean in neurons, but not glial cells of aging brains. In glial cells methylation levels diverged away from the mean, and the tissue-unique signature appears to become stronger. Similarly to neurons, regression to the mean with age was observed in unique sites in CD4+ cells. Future analysis of samples separated by cell type will enable us to determine the relative contribution of each of these mechanisms to epigenetic information loss, as these may differ across organs and pathologies.

Epigenetic information holds diagnostic value. Using the same algorithms and solely relying on methylation data, we were able to classify diseased and healthy organs at high rates. These results highlight the diagnostic potential of methylation data. As some disease induced changes to the methylome are reversible ^7^, and heterochronic parabiosis can reverse or delay some changes to epigenetic age ^8^, it would be interesting to investigate if treatment may reverse the regression to the mean in affected organs. Moreover, it has the potential to suggest which organs are affected by a medical condition. More sophisticated analyses may even distinguish different diseases in the same organ and help classify new disease subtypes. Finally, the epigenetic signature of these tissue-specific sites may allow for their identification from cell-free DNA in blood.

Major limitations of this work include predominantly bulk sample analysis, with limited cell-type and no single-cell data, and lack of longitudinal data. We focused on tissue-level methylation levels across multiple sites. However, each tissue is composed of multiple cell types, often arranged in functionally-important spatial patterns. The bulk analysis limits the extractable information. Cell-type unique methylation sites may have remained undiscovered in our analysis as the signal was suppressed by other cells of that tissue. Similarly, changes that manifest only in a subset of cells may not be detected. For example, beta cells are lost in T2D. However, as they comprise only 1-2% of the pancreas mass, their loss in T2D is unlikely to be captured by this analysis. Indeed, our analysis of pancreas-unique sites identified significant enrichment for pancreas enhancers, however, no enrichment for pancreatic islet enhancers was observed (**Supp. Fig. 3A**). Thus, these experiments do not inform us if epigenetic information loss plays a role in beta cell loss in T2D. The addition of such cell-specific information, both for unique methylation site identification and for deeper tissue analysis, will enable greater sensitivity while informing us which cells are losing information and by which mechanism. Next, while we were able to demonstrate that methylation-based disease classification is possible, our study focused on comparing disease samples to control samples. Future studies should aim to identify changes in methylation patterns within individuals over time, as this would enable us to identify epigenetic information loss in an individual rather than in comparison to a control. Finally, we note that while aggregation of hundreds to thousands of sites across tens of samples enabled robust statistics, the average data content per individual site and patient is low. This is likely the reflection of the stochastic nature of this process.

## Methods

### Unique methylation sites

A site was considered unique for a tissue if the average methylation value in that tissue was either (i) at least 0.1 higher than the 95% quantile of all other tissues (combined) or (ii) at least 0.1 lower than the 5% quantile of all other tissues in the NGDC-CNCB dataset. CD4+ unique sites were identified by applying the same analysis to the NGDC-CNCB blood dataset. Methylation sites with fewer than 2,000 valid values in the dataset were omitted from the analysis. The dataset, containing Illumina HumanMethylation 450 BeadChip probes, was screened for the above criteria using a custom Python script. The results were manually verified for a random subset. Among the 30 available tissues, 8 did not have unique sites. It should be noted that these criteria do not entirely eliminate the possibility of two tissues sharing the same unique site. Parsing the dataset, analyzing the data and sorting the unique sites was done using Python.

### Datasets used

Methylation data were obtained from the Gene Expression Omnibus ^1,65^. All datasets used are listed in **Supplementary Table 2**. Analysis was conducted on the processed Series Matrix Files, which were separated into a methylation data file and an experiment metadata file.

### Motif enrichment

We analyzed the unique sites for de novo and known motifs across all tissues using Hypergeometric Optimization of Motif EnRichment (HOMER) ^66,67^. We extracted ±500bp around the start of methylation probes using the Python Bio.Entrez package ^68^, and obtained chromosome, strands, and genomic coordinate data from the HM450 hg19 database. Sequences were obtained for all unique methylation sites across all tissues and for 3,000 random methylation sites, which were used as background for motif enrichment detection. Sequences were saved in Fasta format using Python. HOMER was run for each tissue’s unique sites separately, against the background sequences, and the motif results were filtered by p-value < 10^-3^ and occurrence percentage in the background <10% using a Python script. Enrichment was calculated as the log2 of the percentage in the target divided by the percentage in the background.

### Chromatin state classification

We analyzed the unique low-in-tissue (blue) using eFORGE 2 (https://eforge.altiusinstitute.org/; ^50^) with the following setting: “--format probeid --bkgd 450k --data erc2-chromatin15state-all –noproxy --reps 1000 --thresh 0.05,0.01”. In cases where the number of unique sites exceeds the capacity of the software (brain), 1,000 sites were randomly selected and used for the analysis.

### Principal component analysis

PCA linear dimensionality reduction was employed to project n-dimensional data (n CpG sites) onto 2D/3D space. We utilized the Python scikit-learn package to perform PCA. The data were parsed and rearranged from a dataframe format to comply with the package’s format requirements, representing probes as dimensions. Whenever a subset of the database was required, we either reconstructed the input files to include only the required dataset or filtered out test subjects in the code prior to implementing PCA on the data. Filtering was conducted on metadata stored in a separate dataframe and/or on methylation sites.

### Disease classification

Values for unique methylation sites were extracted from the databases for each relevant tissue analyzed in the respective datasets and projected onto 2D using PCA. Samples were classified using three models: 1) Ratio of average distances from the sample to healthy and sick samples. 2) Comparing distance from the sample to the control group center of mass and distance from the sample to the disease group center of mass. 3) If the distance from a sample in 2D space was less than twice the standard deviation from the mean location of the disease group, the sample was classified as sick.

To estimate the accuracy of each model, in each iteration, one sample was omitted from the PCA analysis and then transformed and projected onto 2D space using the PCA weights vector. The three models were then applied to each sample, and accuracy was calculated by comparing classification with true results from the database.

All the models projected a single sample onto an existing PCA space based on the rest of the samples, and subsequently classified it into one of the groups without prior knowledge.

### Statistics

P-values were calculated using the binomial distribution in R (dbinom), except when otherwise specified. Where applicable, technical duplicates were removed from data before the analysis.

## Data and materials availability

Generated data are available as supplementary files.

## Acknowledgements

We thank Professors Yosef Gruenbaum, Yuval Ebenstein, Lihi Adler-Abramovich, and Maayan Gal, as well as the Bar lab members, for their critical review and suggestions.

## Author contributions

NS and DZB designed the experiments. NS performed most of the experiments. DZB wrote the manuscript, incorporating edits and comments from NS.

## Data and materials availability

Generated data is available as supplementary files.

## Supplementary Tables

**Supplementary Table 1**. Tissue unique methylation sites, as identified in 23 of the 30 tissues in the NGDC-CNCB dataset.

**Supplementary Table 2**. DNA methylation datasets used.

**Supplementary Table 3**. Tissue-unique methylation data.

**Supplementary Table 4**. CD4+ unique methylation sites, as identified from the NGDC-CNCB blood dataset.

## Supplementary figures

**Supplementary Figure 1.**
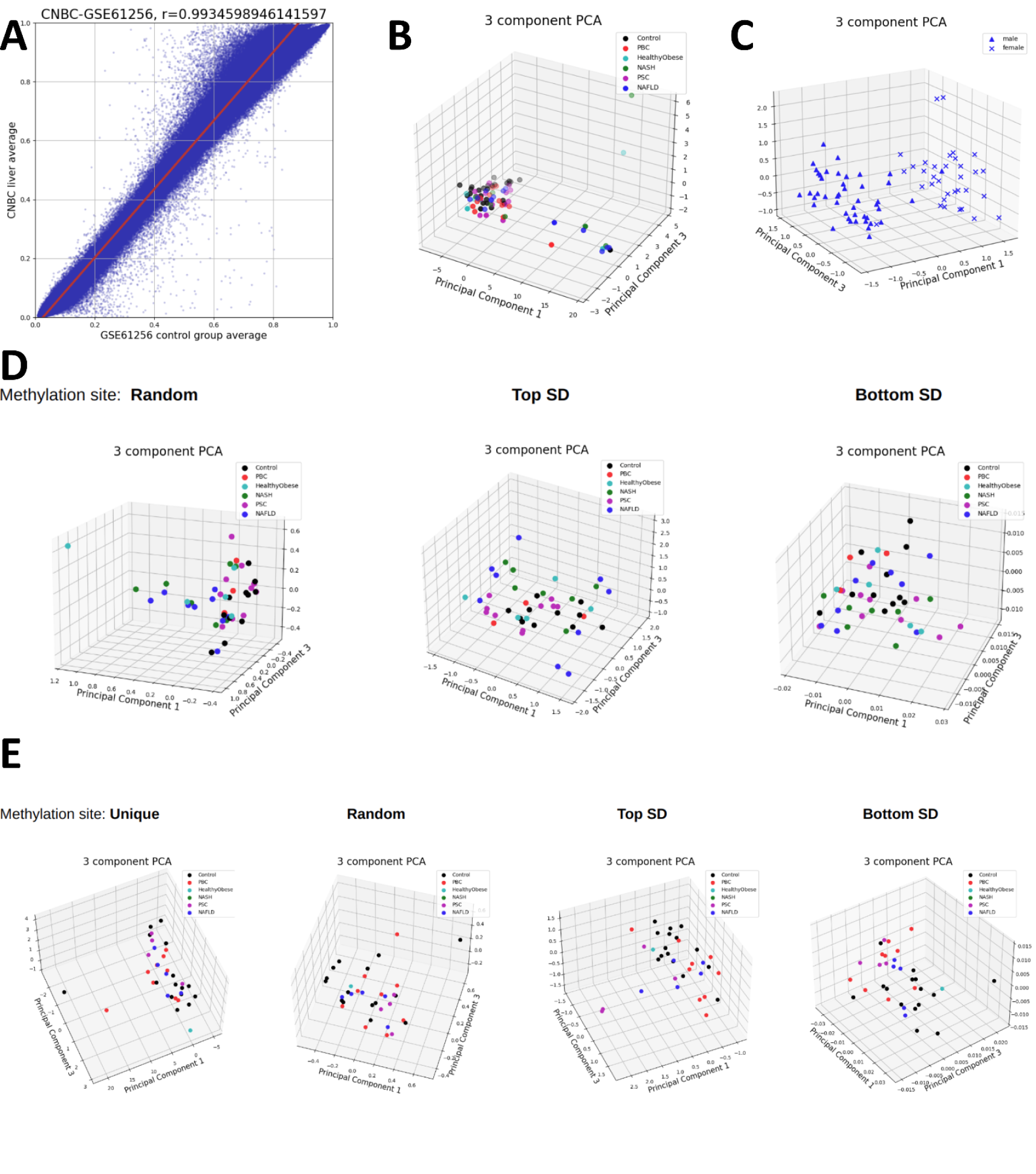
**(A)** Pearson correlation of average liver methylation values in the GSE61256 and the NGDC-CNCB datasets. **(B)** PCA analysis of all GSE61256 liver samples, using uniquely methylated sites. **(C)** PCA analysis of all GSE61256 liver samples using high-variability sites. **(D)** PCA analysis of male liver samples from GSE61256, using random, low-variability and high-variability methylated sites.(**E)** PCA analysis of female liver samples from GSE61256, using unique, random, low-variability and high-variability methylated sites.

**Supplementary Figure 2A.**
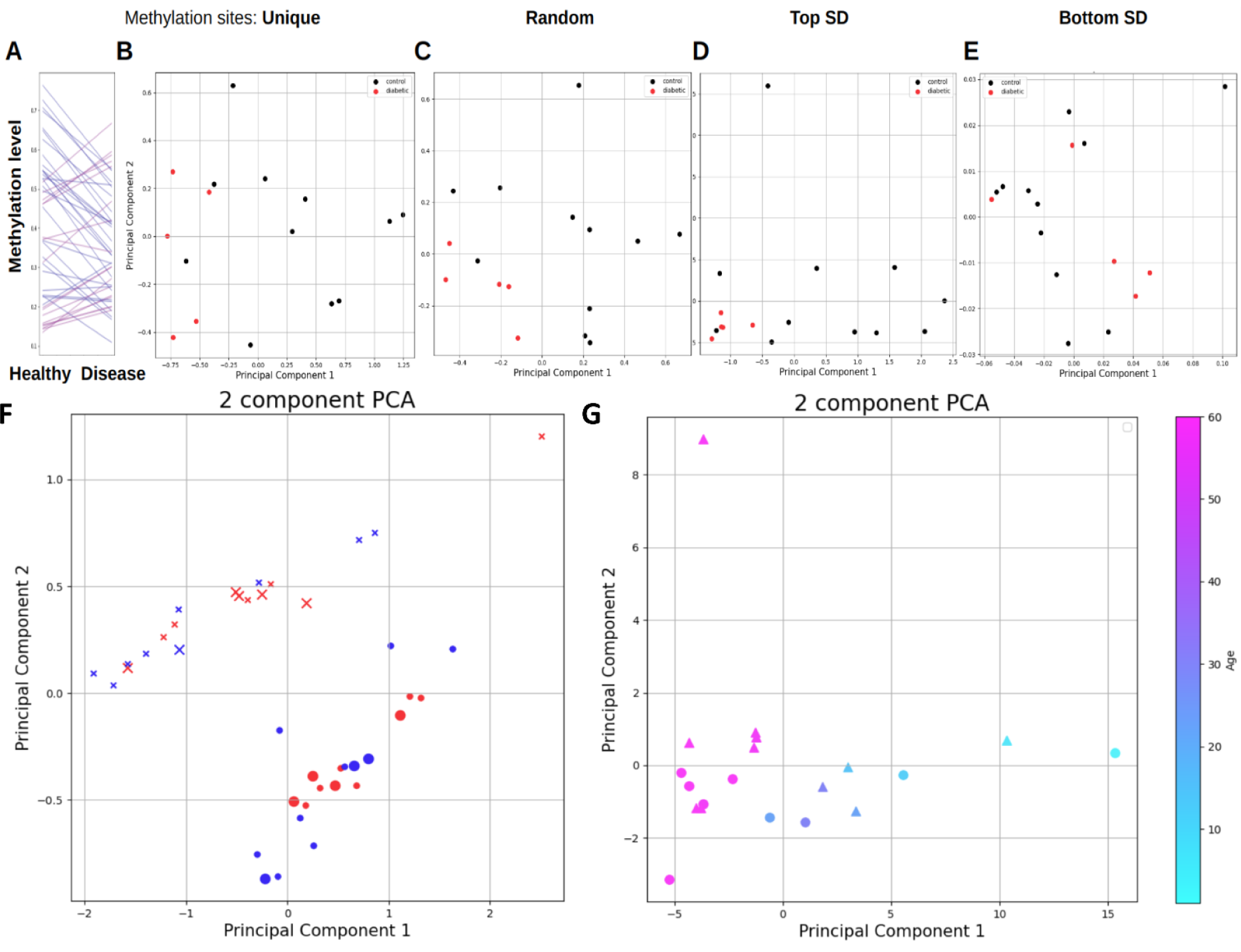
Tissue-unique sites in the pancreas of T2D patients. (**A**) Pancreas-unique methylation sites diverge from the mean both for sites that are more (red) and less (blue) methylated. PCA analysis separates healthy (black) from T2D pancreas when using pancreas-unique (**B**) random (**C**) and high variability (**D**) sites, but not low variability sites (**E**).**(F)** PCA mapping using tissue-unique sites in the adipose of T2D patients. Healthy - blue; diabetic - red. SAT - x; VAT - circle. Size is sex: small - female, large - male.**(G)** PCA analysis of brain samples from GSE38608, using uniquely methylated sites. Circles represent control samples, while triangles represent autism samples. Colors indicate age.

**Supplementary Figure 3.**
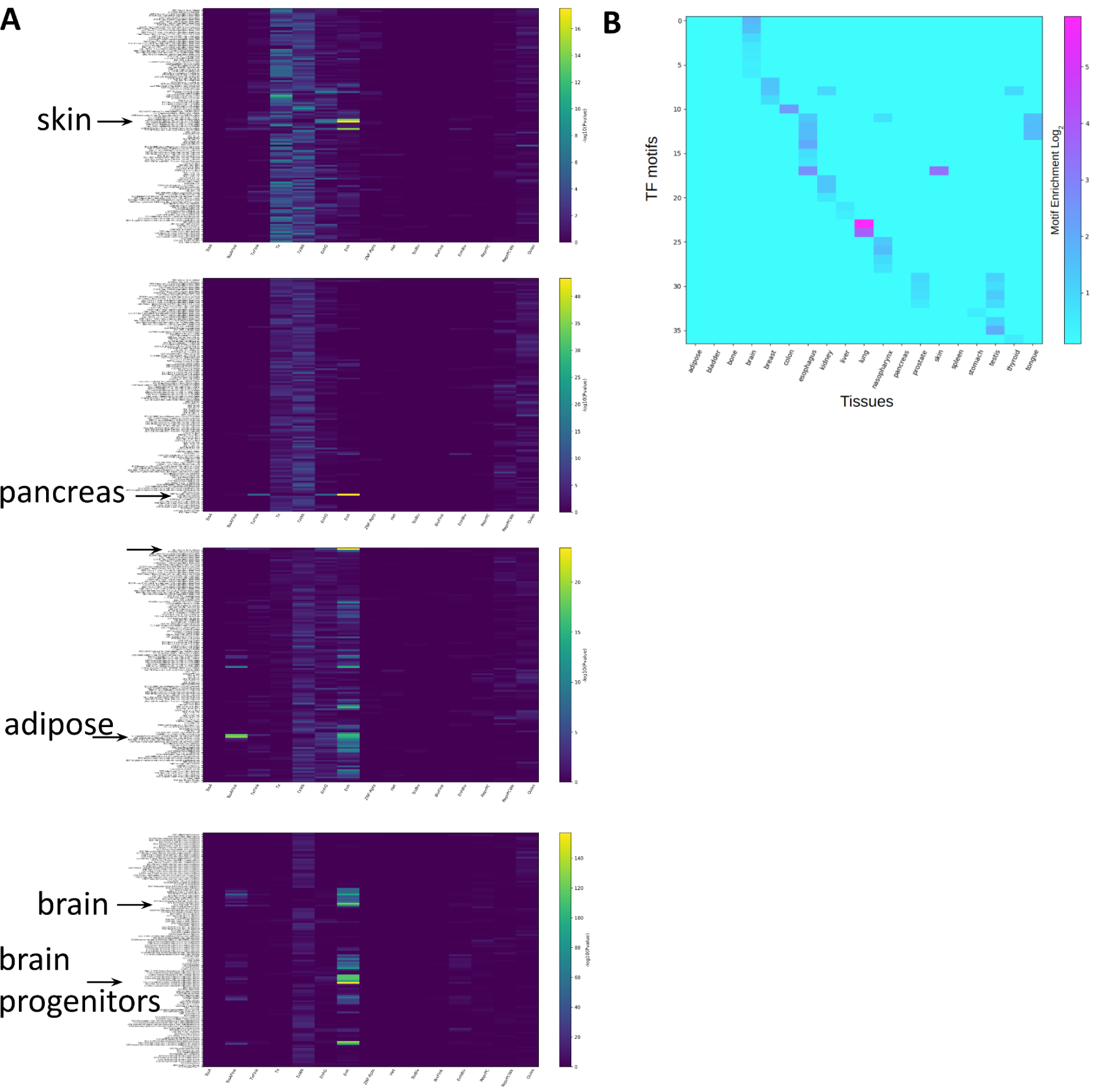
(**A**) Enrichment of unique-low methylation sites across 15 chromatin states (x-axis) and 127 tissues and cell lines (y-axis). Skin, pancreas, adipose and brain are shown in descending order. Arrows and text mark tissues with high enrichment. Full details of chromatin annotation marks can be found at ^51^ (**B**) TF motif enrichment in different tissues.

**Supplementary Figure 4.**
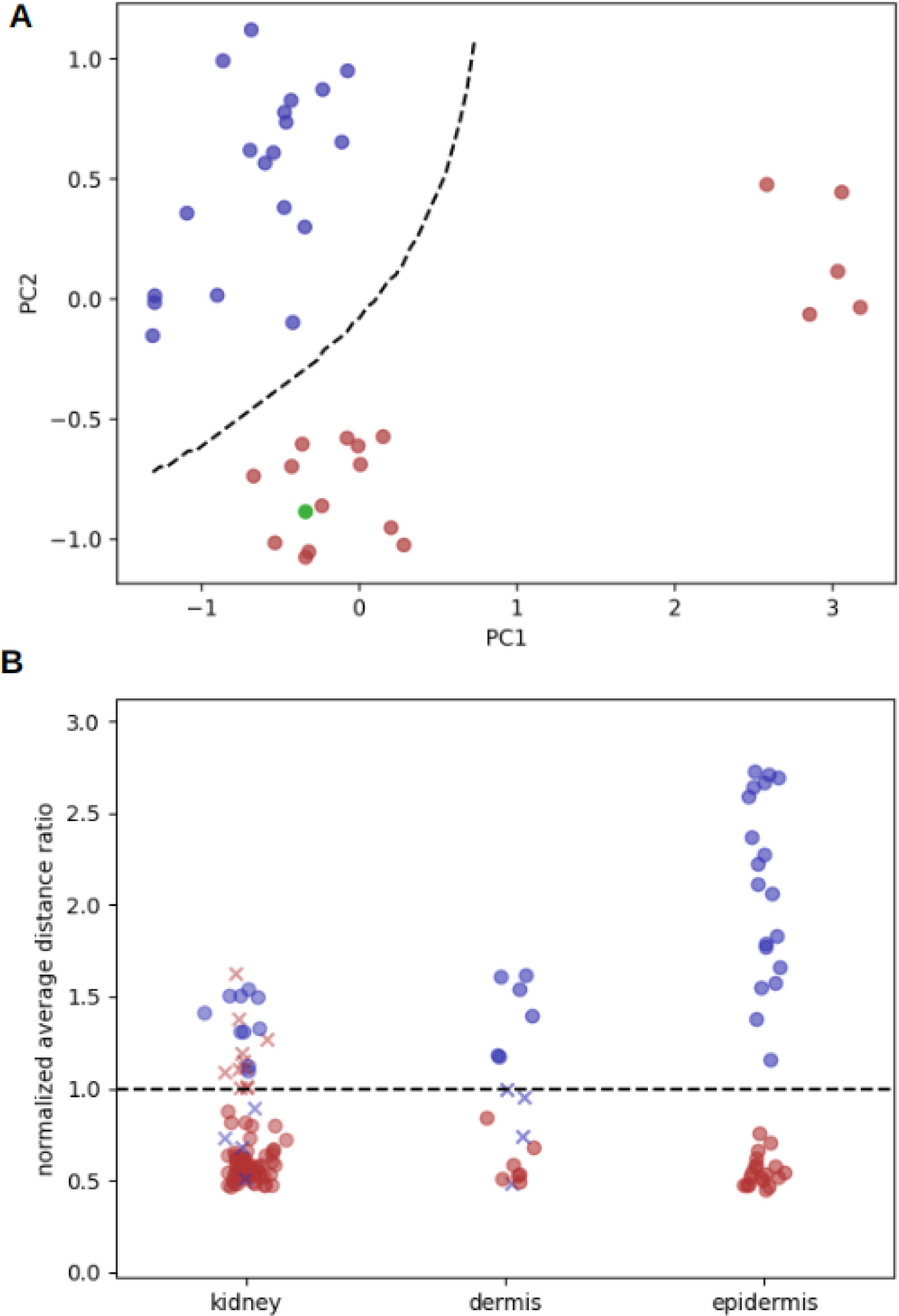
**(A)** Classification of a single epidermis sample (green) in leave-one-out analysis using 2D PCA. The separating (dashed) line was calculated using the formula 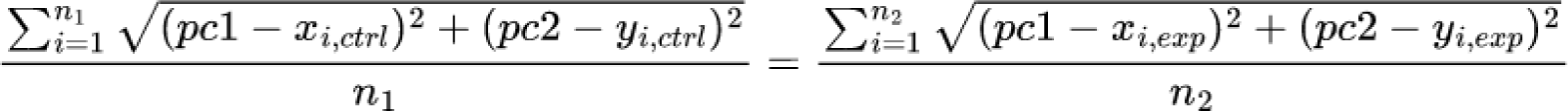 **(B)** Kidney filtration rate and skin exposure classification using PCA and average distance ratio in leave-one-out cross validation analysis. Red dots show high eGFR/ sun exposed skin, while blue dots show low eGFR/ sun protected skin. Correct decisions are marked by full circles whereas incorrect decisions are marked by x.

## References

1. Niccoli, T., and Partridge, L. (2012). Ageing as a risk factor for disease. Curr. Biol. 22, R741–R752.

2. López-Otín, C., Blasco, M.A., Partridge, L., Serrano, M., and Kroemer, G. (2022). Hallmarks of aging: An expanding universe. Cell. 10.1016/j.cell.2022.11.001.

3. López-Otín, C., Blasco, M.A., Partridge, L., Serrano, M., and Kroemer, G. (2013). The hallmarks of aging. Cell 153, 1194–1217.

4. Issa, J.-P. (2014). Aging and epigenetic drift: a vicious cycle. J. Clin. Invest. 124, 24–29.

5. Horvath, S. (2013). DNA methylation age of human tissues and cell types. Genome Biol. 14, R115.

6. Lu, A.T., Quach, A., Wilson, J.G., Reiner, A.P., Aviv, A., Raj, K., Hou, L., Baccarelli, A.A., Li, Y., Stewart, J.D., et al. (2019). DNA methylation GrimAge strongly predicts lifespan and healthspan. Aging 11, 303–327.

7. Poganik, J.R., Zhang, B., Baht, G.S., Tyshkovskiy, A., Deik, A., Kerepesi, C., Yim, S.H., Lu, A.T., Haghani, A., Gong, T., et al. (2023). Biological age is increased by stress and restored upon recovery. Cell Metab. 35, 807–820.e5.

8. Zhang, B., Lee, D.E., Trapp, A., Tyshkovskiy, A., Lu, A.T., Bareja, A., Kerepesi, C., McKay, L.K., Shindyapina, A.V., Dmitriev, S.E., et al. (2023). Multi-omic rejuvenation and life span extension on exposure to youthful circulation. Nat Aging. 10.1038/s43587-023-00451-9.

9. Lu, Y., Brommer, B., Tian, X., Krishnan, A., Meer, M., Wang, C., Vera, D.L., Zeng, Q., Yu, D., Bonkowski, M.S., et al. (2020). Reprogramming to recover youthful epigenetic information and restore vision. Nature 588, 124–129.

10. Yang, J.-H., Griffin, P.T., Vera, D.L., Hayano, M., Meer, M.V., Salfati, E.L., Su, Q., Munding, E.M., Blanchette, M., Bhakta, M., et al. (2019). Erosion of the Epigenetic Landscape and Loss of Cellular Identity as a Cause of Aging in Mammals. bioRxiv, 808642. 10.1101/808642.

11. Sinclair, D.A., and LaPlante, M.D. (2019). Lifespan: Why We Age—and Why We Don’t Have To (Simon and Schuster).

12. Imai, S., and Kitano, H. (1998). Heterochromatin islands and their dynamic reorganization: a hypothesis for three distinctive features of cellular aging. Exp. Gerontol. 33, 555–570.

13. Yang, J.-H., Hayano, M., Griffin, P.T., Amorim, J.A., Bonkowski, M.S., Apostolides, J.K., Salfati, E.L., Blanchette, M., Munding, E.M., Bhakta, M., et al. (2023). Loss of epigenetic information as a cause of mammalian aging. Cell. 10.1016/j.cell.2022.12.027.

14. Bahar, R., Hartmann, C.H., Rodriguez, K.A., Denny, A.D., Busuttil, R.A., Dollé, M.E.T., Calder, R.B., Chisholm, G.B., Pollock, B.H., Klein, C.A., et al. (2006). Increased cell-to-cell variation in gene expression in ageing mouse heart. Nature 441, 1011–1014.

15. Busuttil, R., Bahar, R., and Vijg, J. (2007). Genome dynamics and transcriptional deregulation in aging. Neuroscience 145, 1341–1347.

16. Martinez-Jimenez, C.P., Eling, N., Chen, H.-C., Vallejos, C.A., Kolodziejczyk, A.A., Connor, F., Stojic, L., Rayner, T.F., Stubbington, M.J.T., Teichmann, S.A., et al. (2017). Aging increases cell-to-cell transcriptional variability upon immune stimulation. Science 355, 1433–1436.

17. Warren, L.A., Rossi, D.J., Schiebinger, G.R., Weissman, I.L., Kim, S.K., and Quake, S.R. (2007). Transcriptional instability is not a universal attribute of aging. Aging Cell 6, 775–782.

18. Levy, O., Amit, G., Vaknin, D., Snir, T., Efroni, S., Castaldi, P., Liu, Y.-Y., Cohen, H.Y., and Bashan, A. (2020). Age-related loss of gene-to-gene transcriptional coordination among single cells. Nat Metab 2, 1305–1315.

19. Dritsoula, A., Kislikova, M., Oomatia, A., Webster, A.P., Beck, S., Ponticos, M., Lindsey, B., Norman, J., Wheeler, D.C., Oates, T., et al. (2021). “Epigenome-wide methylation profile of chronic kidney disease-derived arterial DNA uncovers novel pathways in disease-associated cardiovascular pathology.” Epigenetics 16, 718–728.

20. Galbraith, K., and Snuderl, M. (2022). DNA methylation as a diagnostic tool. Acta Neuropathol Commun 10, 71.

21. Cerrato, F., Sparago, A., Ariani, F., Brugnoletti, F., Calzari, L., Coppedè, F., De Luca, A., Gervasini, C., Giardina, E., Gurrieri, F., et al. (2020). DNA Methylation in the Diagnosis of Monogenic Diseases. Genes 11. 10.3390/genes11040355.

22. Sadikovic, B., Levy, M.A., Kerkhof, J., Aref-Eshghi, E., Schenkel, L., Stuart, A., McConkey, H., Henneman, P., Venema, A., Schwartz, C.E., et al. (2021). Clinical epigenomics: genome-wide DNA methylation analysis for the diagnosis of Mendelian disorders. Genet. Med. 23, 1065–1074.

23. Sagy, N., Meyrom, N., Beckerman, P., Pleniceanu, O., and Bar, D.Z. (2022). Kidney-specific methylation patterns correlate with kidney function and are lost upon kidney disease progression. bioRxiv, 2022.09.19.508466. 10.1101/2022.09.19.508466.

24. Xiong, Z., Yang, F., Li, M., Ma, Y., Zhao, W., Wang, G., Li, Z., Zheng, X., Zou, D., Zong, W., et al. (2022). EWAS Open Platform: integrated data, knowledge and toolkit for epigenome-wide association study. Nucleic Acids Res. 50, D1004–D1009.

25. Jeong, E.H., Jun, D.W., Cho, Y.K., Choe, Y.G., Ryu, S., Lee, S.M., and Jang, E.C. (2013). Regional prevalence of non-alcoholic fatty liver disease in Seoul and Gyeonggi-do, Korea. Clin. Mol. Hepatol. 19, 266–272.

26. Amarapurkar, D., Kamani, P., Patel, N., Gupte, P., Kumar, P., Agal, S., Baijal, R., Lala, S., Chaudhary, D., and Deshpande, A. (2007). Prevalence of non-alcoholic fatty liver disease: population based study. Ann. Hepatol. 6, 161–163.

27. Kim, I.H., Kisseleva, T., and Brenner, D.A. (2015). Aging and liver disease. Curr. Opin. Gastroenterol. 31, 184–191.

28. Horvath, S., Erhart, W., Brosch, M., Ammerpohl, O., von Schönfels, W., Ahrens, M., Heits, N., Bell, J.T., Tsai, P.-C., Spector, T.D., et al. (2014). Obesity accelerates epigenetic aging of human liver. Proc. Natl. Acad. Sci. U. S. A. 111, 15538–15543.

29. García-Calzón, S., Perfilyev, A., de Mello, V.D., Pihlajamäki, J., and Ling, C. (2018). Sex Differences in the Methylome and Transcriptome of the Human Liver and Circulating HDL-Cholesterol Levels. J. Clin. Endocrinol. Metab. 103, 4395–4408.

30. Zhang, Y., Klein, K., Sugathan, A., Nassery, N., Dombkowski, A., Zanger, U.M., and Waxman, D.J. (2011). Transcriptional profiling of human liver identifies sex-biased genes associated with polygenic dyslipidemia and coronary artery disease. PLoS One 6, e23506.

31. Lau-Corona, D., Bae, W.K., Hennighausen, L., and Waxman, D.J. (2020). Sex-biased genetic programs in liver metabolism and liver fibrosis are controlled by EZH1 and EZH2. PLoS Genet. 16, e1008796.

32. Clodfelter, K.H., Holloway, M.G., Hodor, P., Park, S.-H., Ray, W.J., and Waxman, D.J. (2006). Sex-dependent liver gene expression is extensive and largely dependent upon signal transducer and activator of transcription 5b (STAT5b): STAT5b-dependent activation of male genes and repression of female genes revealed by microarray analysis. Mol. Endocrinol. 20, 1333–1351.

33. Kasarinaite, A., Sinton, M., Saunders, P.T.K., and Hay, D.C. (2023). The Influence of Sex Hormones in Liver Function and Disease. Cells 12. 10.3390/cells12121604.

34. Bogardus, C., Lillioja, S., Howard, B.V., Reaven, G., and Mott, D. (1984). Relationships between insulin secretion, insulin action, and fasting plasma glucose concentration in nondiabetic and noninsulin-dependent diabetic subjects. J. Clin. Invest. 74, 1238–1246.

35. Reaven, G.M. (1988). Banting lecture 1988. Role of insulin resistance in human disease. Diabetes 37, 1595–1607.

36. Macauley, M., Percival, K., Thelwall, P.E., Hollingsworth, K.G., and Taylor, R. (2015). Altered volume, morphology and composition of the pancreas in type 2 diabetes. PLoS One 10, e0126825.

37. Bahour, N., Cortez, B., Pan, H., Shah, H., Doria, A., and Aguayo-Mazzucato, C. (2022). Diabetes mellitus correlates with increased biological age as indicated by clinical biomarkers. Geroscience 44, 415–427.

38. Volkmar, M., Dedeurwaerder, S., Cunha, D.A., Ndlovu, M.N., Defrance, M., Deplus, R., Calonne, E., Volkmar, U., Igoillo-Esteve, M., Naamane, N., et al. (2012). DNA methylation profiling identifies epigenetic dysregulation in pancreatic islets from type 2 diabetic patients. EMBO J. 31, 1405–1426.

39. Bergman, R.N., Kim, S.P., Catalano, K.J., Hsu, I.R., Chiu, J.D., Kabir, M., Hucking, K., and Ader, M. (2006). Why visceral fat is bad: mechanisms of the metabolic syndrome. Obesity 14 *Suppl 1*, 16S – 19S.

40. Jin, N., Lee, H.-M., Hou, Y., Yu, A.C.S., Li, J.-W., Kong, A.P.S., Lam, C.C.H., Wong, S.K.H., Ng, E.K.W., Ma, R.C.W., et al. (2021). Integratome analysis of adipose tissues reveals abnormal epigenetic regulation of adipogenesis, inflammation, and insulin signaling in obese individuals with type 2 diabetes. Clin. Transl. Med. 11, e596.

41. Ginsberg, M.R., Rubin, R.A., Falcone, T., Ting, A.H., and Natowicz, M.R. (2012). Brain transcriptional and epigenetic associations with autism. PLoS One 7, e44736.

42. Rittié, L., and Fisher, G.J. (2015). Natural and sun-induced aging of human skin. Cold Spring Harb. Perspect. Med. 5, a015370.

43. Debacq-Chainiaux, F., Leduc, C., Verbeke, A., and Toussaint, O. (2012). UV, stress and aging. Dermatoendocrinol. 4, 236–240.

44. Vandiver, A.R., Irizarry, R.A., Hansen, K.D., Garza, L.A., Runarsson, A., Li, X., Chien, A.L., Wang, T.S., Leung, S.G., Kang, S., et al. (2015). Age and sun exposure-related widespread genomic blocks of hypomethylation in nonmalignant skin. Genome Biol. 16, 80.

45. Horvath, S., and Raj, K. (2018). DNA methylation-based biomarkers and the epigenetic clock theory of ageing. Nat. Rev. Genet. 19, 371–384.

46. Muñoz-Najar, U., and Sedivy, J.M. (2011). Epigenetic control of aging. Antioxid. Redox Signal. 14, 241–259.

47. Funk, M.C., Zhou, J., and Boutros, M. (2020). Ageing, metabolism and the intestine. EMBO Rep. 21, e50047.

48. Noreen, F., Röösli, M., Gaj, P., Pietrzak, J., Weis, S., Urfer, P., Regula, J., Schär, P., and Truninger, K. (2014). Modulation of age- and cancer-associated DNA methylation change in the healthy colon by aspirin and lifestyle. J. Natl. Cancer Inst. 106. 10.1093/jnci/dju161.

49. Bell, C.G., Lowe, R., Adams, P.D., Baccarelli, A.A., Beck, S., Bell, J.T., Christensen, B.C., Gladyshev, V.N., Heijmans, B.T., Horvath, S., et al. (2019). DNA methylation aging clocks: challenges and recommendations. Genome Biol. 20, 249.

50. Breeze, C.E., Paul, D.S., van Dongen, J., Butcher, L.M., Ambrose, J.C., Barrett, J.E., Lowe, R., Rakyan, V.K., Iotchkova, V., Frontini, M., et al. (2016). eFORGE: A Tool for Identifying Cell Type-Specific Signal in Epigenomic Data. Cell Rep. 17, 2137–2150.

51. Roadmap Epigenomics Consortium, Kundaje, A., Meuleman, W., Ernst, J., Bilenky, M., Yen, A., Heravi-Moussavi, A., Kheradpour, P., Zhang, Z., Wang, J., et al. (2015). Integrative analysis of 111 reference human epigenomes. Nature 518, 317–330.

52. Guintivano, J., Aryee, M.J., and Kaminsky, Z.A. (2013). A cell epigenotype specific model for the correction of brain cellular heterogeneity bias and its application to age, brain region and major depression. Epigenetics 8, 290–302.

53. Download https://ngdc.cncb.ac.cn/ewas/datahub/download.

54. Cutler, R.G. (1982). The dysdifferentiative hypothesis of mammalian aging and longevity. The aging brain 20, 1–18.

55. Fu, Y., Yu, Y., Paxinos, G., Watson, C., and Rusznák, Z. (2015). Aging-dependent changes in the cellular composition of the mouse brain and spinal cord. Neuroscience 290, 406–420.

56. Wang, Q., Qi, Y., Shen, W., Xu, J., Wang, L., Chen, S., Hou, T., and Si, J. (2021). The Aged Intestine: Performance and Rejuvenation. Aging Dis. 12, 1693–1712.

57. Lee, H., Hong, Y., and Kim, M. (2021). Structural and Functional Changes and Possible Molecular Mechanisms in Aged Skin. Int. J. Mol. Sci. 22. 10.3390/ijms222212489.

58. Csekes, E., and Račková, L. (2021). Skin Aging, Cellular Senescence and Natural Polyphenols. Int. J. Mol. Sci. 22. 10.3390/ijms222312641.

59. Kalluri, R., and Weinberg, R.A. (2009). The basics of epithelial-mesenchymal transition. J. Clin. Invest. 119, 1420–1428.

60. Roeder, S.S., Stefanska, A., Eng, D.G., Kaverina, N., Sunseri, M.W., McNicholas, B.A., Rabinovitch, P., Engel, F.B., Daniel, C., Amann, K., et al. (2015). Changes in glomerular parietal epithelial cells in mouse kidneys with advanced age. Am. J. Physiol. Renal Physiol. 309, F164–F178.

61. Zeisberg, M., Yang, C., Martino, M., Duncan, M.B., Rieder, F., Tanjore, H., and Kalluri, R. (2007). Fibroblasts derive from hepatocytes in liver fibrosis via epithelial to mesenchymal transition. J. Biol. Chem. 282, 23337–23347.

62. Kim, K.K., Kugler, M.C., Wolters, P.J., Robillard, L., Galvez, M.G., Brumwell, A.N., Sheppard, D., and Chapman, H.A. (2006). Alveolar epithelial cell mesenchymal transition develops in vivo during pulmonary fibrosis and is regulated by the extracellular matrix. Proc. Natl. Acad. Sci. U. S. A. 103, 13180–13185.

63. Bechtel, W., McGoohan, S., Zeisberg, E.M., Müller, G.A., Kalbacher, H., Salant, D.J., Müller, C.A., Kalluri, R., and Zeisberg, M. (2010). Methylation determines fibroblast activation and fibrogenesis in the kidney. Nat. Med. 16, 544–550.

64. Galle, E., Thienpont, B., Cappuyns, S., Venken, T., Busschaert, P., Van Haele, M., Van Cutsem, E., Roskams, T., van Pelt, J., Verslype, C., et al. (2020). DNA methylation-driven EMT is a common mechanism of resistance to various therapeutic agents in cancer. Clin. Epigenetics 12, 27.

65. Barrett, T., Wilhite, S.E., Ledoux, P., Evangelista, C., Kim, I.F., Tomashevsky, M., Marshall, K.A., Phillippy, K.H., Sherman, P.M., Holko, M., et al. (2013). NCBI GEO: archive for functional genomics data sets--update. Nucleic Acids Res. 41, D991–D995.

66. Duttke, S.H., Chang, M.W., Heinz, S., and Benner, C. (2019). Identification and dynamic quantification of regulatory elements using total RNA. Genome Res. 29, 1836–1846.

67. Heinz, S., Benner, C., Spann, N., Bertolino, E., Lin, Y.C., Laslo, P., Cheng, J.X., Murre, C., Singh, H., and Glass, C.K. (2010). Simple combinations of lineage-determining transcription factors prime cis-regulatory elements required for macrophage and B cell identities. Mol. Cell 38, 576–589.

68. Cock, P.J.A., Antao, T., Chang, J.T., Chapman, B.A., Cox, C.J., Dalke, A., Friedberg, I., Hamelryck, T., Kauff, F., Wilczynski, B., et al. (2009). Biopython: freely available Python tools for computational molecular biology and bioinformatics. Bioinformatics 25, 1422–1423.

